# The effect of different exercise training modes on dentate gyrus neurodegeneration and synaptic plasticity in morphine-dependent rats

**DOI:** 10.1101/2021.04.13.439604

**Authors:** Kamal Ranjbar, Ebrahim Zarrinkalam, Sara Soleimani Asl, Iraj Salehi, Masoumeh Taheri, Alireza Komaki

**Author notes:** Corresponding author: Prof. Alireza Komaki, Neurophysiology Research Center, Hamadan University of Medical Sciences, Hamadan, Iran., E-mail addresses, URL: http://umsha.ac.ir (A. Komaki). Second Corresponding author: Dr. Ebrahim Zarrinkalam, Department of Physical Education, Faculty of Physical Education and Sport Sciences, Islamic Azad University, Hamedan Branch, Hamedan, Iran. **Data Availability Statement**: All relevant data are within the manuscript and its Supporting Information files. **Competing interests**: The authors have declared that no competing interests exist.

## Abstract

Various impacts of exercise on brain performance have been documented following morphine dependence induction; however, the underlying neuronal mechanisms remain unclear. The present research was done to investigate the impact of different exercise training modes on neuronal maturation, and synaptic plasticity in the perforant pathway (PP)-dentate gyrus (DG) synapse in the morphine-dependent rats. Five groups, including a control group (Con, ten healthy rats) and forty morphine-dependent rats were considered as follows (n=10/group): 1) sedentary-dependent (Sed-D); 2) endurance exercise-dependent (En-D); 3) strength exercise-dependent (St-D); and 4) concurrent exercise-dependent (Co-D). The exercise training groups were subjected to endurance, strength, and concurrent training 5 days per week for 10 weeks. After training sessions, the field excitatory postsynaptic potential (fEPSP) slope and population spike (PS) amplitude in DG were determined in response to high-frequency stimulation (HFS) of the PP. For assessing neurogenesis NeuroD level was evaluated after performing all experiments. Concurrent training increased PS amplitude and EPSP than the control group. NeuroD in the morphine-dependent rats significantly decreased, but concurrent training returned the NeuroD to the healthy rat level. Concurrent training can ameliorate synaptic plasticity impairment in morphine-dependent rats through neurogenesis promotion. According to the results, concurrent training can be an appropriate novel candidate for treating opioid addiction.

## Introduction

It has been shown that morphine addiction reduces learning and memory abilities and disrupts dentate gyrus (DG) synaptic plasticity via changes in synaptic transmission effectiveness [1-3]. In this regard, several studies have recently indicated that acute morphine exposure can modify the nervous system [4, 5], and hippocampal synaptic plasticity retrogression can be observed in those addicted to morphine [6]. It should be noted that repeated exposure to opioids can induce oxidative stress-related neurodegeneration, apoptosis [7], and neurogenesis [8], as well as a reduction in expression levels of postsynaptic density protein 95 (PSD-95), leading to a decrease in synapse efficacy and hippocampus function and impairment of the related behaviors [9]. However, the underlying neuronal mechanisms of learning and memory deficit following morphine dependence remain unknown [3].

Long-term potentiation (LTP) has been introduced as an experimental model to assess activity-dependent synaptic plasticity and also has been known as a cellular mechanism for learning and memory formation [3]. It has been suggested that morphine dependence impairs LTP induction in the rat hippocampal region [6, 10, 11]. Reversing or preventing LTP reduction following morphine dependence may ameliorate brain function.

Regular exercise training is an alternative therapeutic strategy to reverse or prevent pathological changes observed in morphine-dependent rats. Exercise training increase memory function in humans [12, 13] and animals [14]. It is beneficial for the hippocampus and attenuates memory impairment in morphine-dependent rats [15]; however, a few studies have been conducted on the type of exercise training. In this regard, Lin et al. demonstrated that various kinds of exercise can exert differential impacts on memory and neuronal compatibility in several brain areas [16]. Our previous study showed the effectiveness of different kinds of exercise on learning and memory function [17]. However, it is still unknown why different types of exercise can result in various impacts on brain function following induction of morphine dependence and the most effective exercise type (endurance, strength, or concurrent (the combination of both endurance and resistance)) on LTP amelioration has not yet been found in morphine-dependent rats.

Neurogenesis is the main determinant of synaptic plasticity; thus, this study aimed at investigating the impacts of different exercise training modes on neurogenesis, and LTP in the perforant pathway (PP)-DG granule cell synapse of the morphine-dependent rats. We also tried to identify the most effective type of exercise training in LTP impairments due to morphine injection in male rats.

## Materials and Method

### Animals

Fifty 6-8-week old adult male Wistar rats weighting 150-180 g were kept in polypropylene cages (50 × 26 × 25 cm) (five per cage) in an air-conditioned room (21-23°C) with 12-h light-dark cycle (lights on from 08:00 to 20:00) at 55 ± 3% humidity. Food and water were freely accessible. The rats were purchased from the Hamadan University of Medical Sciences, Iran. All experiments were conducted based on the National Institutes of Health and the Guide for the Care and Use of Laboratory Animals (NIH Publication 80-23, 1996). The Ethical Committee of Hamadan University of Medical Sciences confirmed the research protocol.

### Morphine Dependence Induction

Initially, drinking water was added to 0.1 mg/mL of morphine sulfate (Temad Company, Iran) for two consecutive days; 0.2 mg/mL was added on the third and fourth days, 0.3 mg/mL on the fifth and sixth days, and 0.4 mg/mL on the seventh to twenty-first days. Sucrose (40 mg/mL) was added to eliminate the bitter taste of the morphine sulfate in the first two weeks of the research [17]. It should be noted that the control group received water (30-50 mL/rat) without sucrose.

### Withdrawal rating scale

To examine morphine dependence, 1 mg/kg of naloxone hydrochloride (Sigma-Aldrich, St. Louis, MO, USA) was administered via intraperitoneal (IP) injection one day after morphine treatment. After the injection of naloxone, the rats were placed in a specific quiet room and their behavior was noted for 30 min through a rater blinded to the study protocol and their withdrawal signs (wet-dog shakes, genital licks, circles, rearing, and jumping) were monitored and scored based on the modified version of Gellert-Holtzman scale [18].

### Experimental design

When morphine dependence was confirmed by the naloxone test, a day later, the animals were assigned into five groups (10 animals per group): 1) sedentary-dependent (Sed-D), 2) endurance exercise-dependent (En-D), strength exercise-dependent (St-D), concurrent exercise dependent (Co-D), and 5) control (healthy; Con) groups. The training schedule was performed in two stages: adaptation and exercise. Adaptation (running on the treadmill for aerobic training and climbing up from a ladder in resistance training) lasted for one week (3 sessions, each session lasted 10 min). Figure 1 illustrates the experimental timeline and Table 1 presents the general features of the experimental groups.

**Table-1:** General characteristics of the experimental groups

| **Group** | **Con** | **Sed-D** | **En-D** | **St-D** | **Co-D** |
| --- | --- | --- | --- | --- | --- |
| **Number** | 10 | 10 | 10 | 10 | 10 |
| **Age (Week)** | 6-8 | 6-8 | 6-8 | 6-8 | 6-8 |
| **Initial body mass (g)** | 164±11 | 158±07 | 172±12 | 160±08 | 172±15 |
| **Final body mass (g)** | 339±24 | 345±31 | 336±28 | 352±38 | 340±27 |

Co-D: concurrent training-dependent group, St-D, strength training-dependent group, Sed-D: sedentary morphine-dependent group, Con: control group, En-D: endurance training-dependent group.

**Figure 1.**
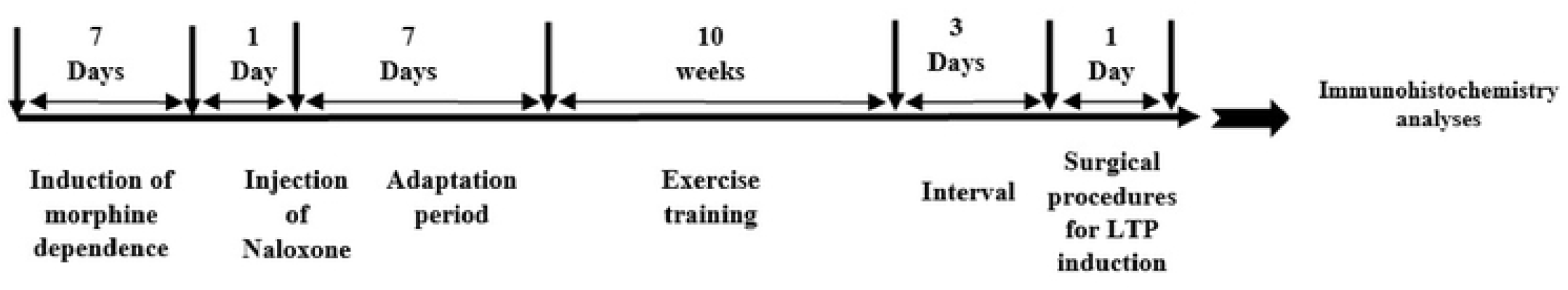
Experimental timeline. After morphine dependence induction, the exercise training groups were subjected to the 10-week (5 days a week) endurance, strength, and concurrent training. At the end of the training, the rats were subjected to urethane anesthesia (IP injection) and located in a stereotaxic device for surgical procedures and electrophysiological recording. The field excitatory postsynaptic potential (fEPSP) slope and population spike (PS) amplitude in the dentate gyrus (DG) were determined after perforant pathway (PP) stimulation. After obtaining a stable baseline of responses for at least 20 min, long-term potentiation (LTP) was induced via the high-frequency stimulation protocol. Finally, for neurogenesis NeuroD level were assessed.

### Exercise training

#### Endurance training

Endurance exercise-dependent rats performed endurance training. After the adaptation stage, the subjects in the aerobic training groups were subjected to running on a treadmill with eight individual lanes for ten weeks (5 days per week). Intensity and duration of the exercise gradually increased from the first week of training (10m/min/5° incline/10 min) to the tenth week (30 m/min/12° incline/60 min).

#### Resistance training

Strength exercise-dependent group rats performed resistance training that was started 3 days after adaptation. Resistance training was conducted according to the Philippe et al. method [19] after some changes. In this group, the rats were trained to climb a 1 m high ladder at an 85° incline, 12 times per session (5 days per week) for 10 weeks. Animals were located at the bottom of ladder and forced for ladder climbing using tweezers (tweezers were used from primitive sessions), while the weights in a cloth bag were hung up to their tails. It should be noted that after a few sessions, the rats instinctively climbed up the ladder without any tension. In the beginning, the weights were 50% of the rats’ body mass followed by a gradual increase to 130% within 10 weeks (1st and 2nd weeks: 50-60%; 3rd to 5th weeks: 70-90%; 6th to 8th weeks: 100-110%; and 9th to 10th weeks: 110-130%). The rats were weighed every two weeks. The exercise sessions included three sets of four repetitions with an interval of 15 s and 3 min during repetitions and sets, respectively. The rats could rest when they reached the top in a compartment on the top of the ladder.

#### Concurrent training

Concurrent exercise-dependent rats were subjected to the combined resistance-endurance training based on our recently published protocol [17] in half of each session. The climbing procedure included two sets of three repetitions followed by running on a treadmill for 30 min (30 m/min) [20]. Similar to the other two training groups, the intensity and duration of the exercise were gradually increased.

### Surgical procedures

The methodologies used for surgeries were similar to previous studies published by our laboratory [21-23]. In brief, anesthetization was done 3 days after the last exercise session, using urethane administration intraperitoneally (1.5 g/kg, Sigma-Aldrich, USA, supplemental administrations as needed). For surgical procedures, implanting electrodes, and field potential recording, the animal’s head was fixed in the stereotaxic device (Stoelting Co., Wood Dale, IL, USA). For maintaining the rat’s body temperature at 36.5 ± 0.5°C, a heating pad, which should be electrically shielded was employed during the experimental procedures. Small burr holes were made in the skull with a drill after making the initial skin incision. In the next step, implantation of the two bipolar Teflon-coated stainless steel electrodes (not for tips) with a diameter of 125 mm into the skull was done (Advent Co., UK). Electrode positioning was as follows [24]: placing the stimulating electrode into the PP (coordinates: 8.1 mm posterior to the bregma, 4.3 mm lateral to the midline, and 3.2 mm ventral below the skull) and the recording electrode into the granule cells of the DG (coordinates: 3.8 mm posterior to bregma and, 2.3 mm lateral to the midline) after observing the maximal field excitatory postsynaptic potentials (fEPSPs) (commonly 2.7–3.2 mm ventral). For preventing brain tissue injuries, we lowered the electrodes very gradually (0.2 mm/min) from cortex to hippocampus.

Through the electrophysiological monitoring system, accurate ventral electrode positioning was achieved, in which the DG-evoked responses after a single-pulse PP stimulation are monitored.

### Electrophysiological recording and hippocampal LTP recording protocol

As described in our previous studies [21, 25, 26], the current input/output specifications were determined via PP stimulation for calculating the stimulus intensity for all rats (40% maximal population spike (PS)). Accordingly, a single pulse of 0.1 ms (0.1 Hz) was given using a constant-current stimulus isolation unit (A365; WPI, USA). Granular cells of the DG stimulated by PP were used for recording field potential. PP was given the test stimuli per 10 s. The positioning of the electrodes was for obtaining the greater PS and fEPSP amplitude. High-frequency stimulation (HFS) protocol was applied for LTP induction after a steady-state baseline response (about 40 min) using the following characteristics: 10 bursts of 20 stimuli at 400 Hz, the length of 0.2 ms, and an interburst interval of 10 s. The HSF was adjusted at the stimulus intensity to evoke PS amplitude and fEPSP slope of about 80% of the maximum response. For determining the alterations in the synaptic response of DG cells, fEPSP, as well as PS, were noted 5, 30, and 60 min after HFS. Ten continuous evoked responses were averaged with an interval of 10 s at each time point.

Using homemade software, the stimuli characteristics were determined (eTrace, www.sciencebeam.com). They were transferred through a data acquisition board connected to the A365 constant-current isolator unit before giving to the PP by stimulus electrodes. The resulted field potential response arisen from DG was moved through a preamplifier. It was then amplified (1000×) (Differential amplifier DAM 80 WPI, USA) and filtered (bandpass: 1-3 kHz). This was also digitized at 10 kHz and could be observed on a computer. The obtained values were recorded in a file for facilitating further offline analysis.

### Immunohistochemistry

Neuronal differentiation factor NeuroD (also known as Beta2) is a proneural basic helix-loop-helix (bHLH) transcription factor involved in the increased neuronal survival, proliferation, and differentiation to neural progenitor cells in the DG of the hippocampus [27]. Neurogenesis includes three stages: cell differentiation, proliferation, and survival [28]. It is well established that NeuroD is involved in embryonic and postnatal neurogenesis [29]. For determining NeuroD-positive cells, briefly, intracardial perfusion of the rat’s brain (5 rats in each group) was first carried out by PBS, and then 4% paraformaldehyde in fetal bovine serum (FBS) (0.1 M, pH 7.4). The animal brain was quickly removed and fixed by immersion in 4% paraformaldehyde at 4°C for 72 h. Its materials were dehydrated in ascending alcohol series, and after xylene, baptist wax, and paraffin embedding, using MICROM HM 500 OM microtome, the hippocampal tissue blocks were separated into 5 μm sections. They were then washed three times within 5 min via 1xTBS pre-incubation using a blocking solution composed of 10% normal serum and 0.25% Triton X-100 in PBS at room temperature (1 h). After deparaffinization and rehydration, the sections were immunostained by goat polyclonal anti-NeuroD (Santa Cruz Biotechnology, Santa Cruz, CA, USA, 1:200) to visualize NeuroD-positive cells. The sections then washed three times again in 5 min with 1x TBS, followed by exposure to corresponding biotinylated secondary antibody (rabbit anti-goat IgG) at a dilution of 1: 200 (Vector) and Cy3-conjugated streptavidin at a dilution of 1: 100 (Sigma) within 60 min at 25°C. In the next stage, they were subjected to washing in 1x TBS and mounted on slides. The number of the dark brown cells expressing NeuroD in the hippocampal DG was counted using the Motic Advanced Plus 2 software after taking digitized images by Olympus BX53 microscope (Shinjuku, Tokyo, Japan), in three to five different fields per section at an absolute magnification of 400X, and five rats per group were examined.

### Histology

At the end of the experiments, using brain sections, the stimulating and recording areas in the hippocampus were histologically approved. In the final stage, anesthesia was induced by urethane followed by perfusion through formol saline [30]. The samples were cut into 50 μm thick serial sections and were subjected to Hematoxylin-Eosin (H&E) staining for histological confirmation and also the validation of the position of the electrode tip [21, 31] (Fig. 2).

**Figure 2.**
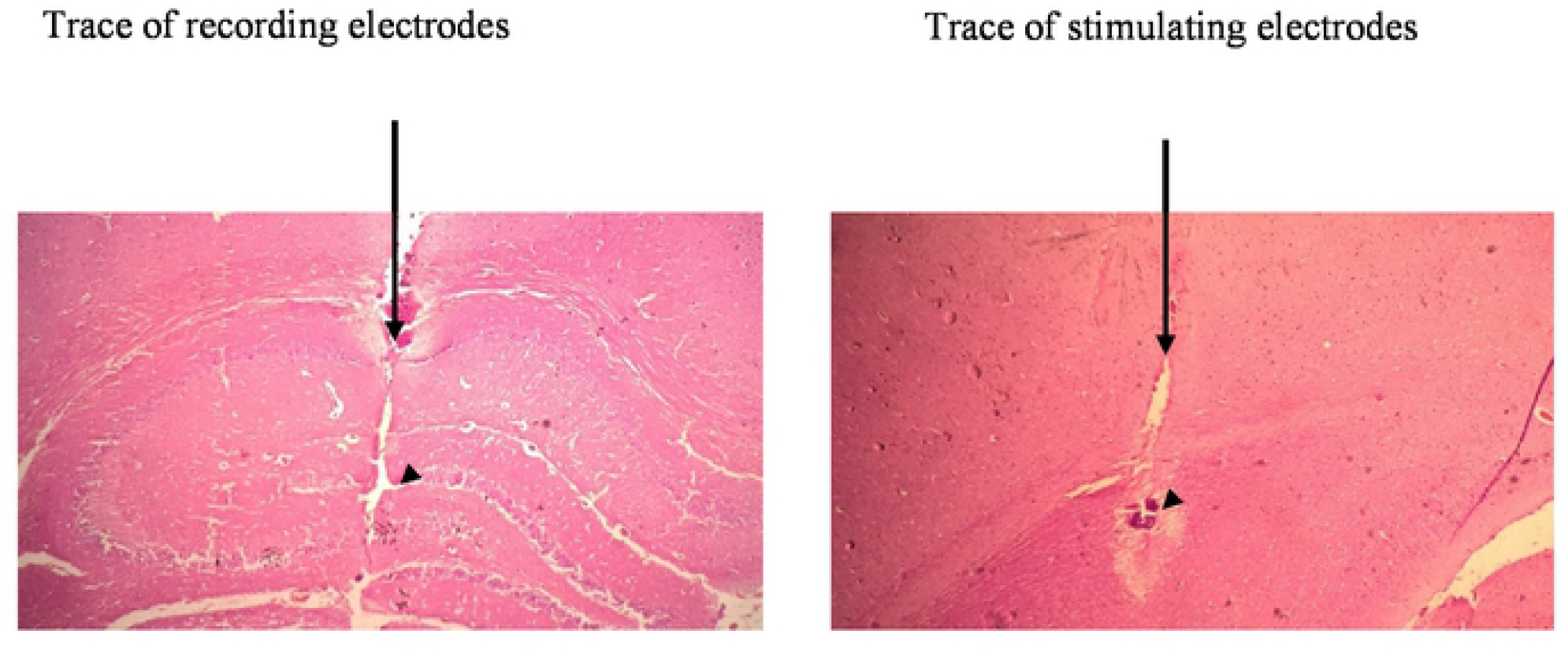
Exemplary photomicrograph indicating the positions of stimulating and recording electrodes tips (arrowheads) in a hippocampus sagittal section. Stimulating and recording electrode traces are observed in the right and the left sides (arrows), respectively. Scale bar: 0.5 mm.

### Statistical analysis

Data analysis was performed by the GraphPadPrism® 5.0 software (GraphPad Software Inc, USA). Data are shown as the mean ± standard error of the mean (SEM). One-way analysis of variance (ANOVA) and 5×4 (group×time) mixed-plot factorial repeated-measures ANOVA were applied to analyze the data. We employed the Tukey post-hoc test to compare the experimental groups. P-values of less than 0.05 were regarded as significant.

## Results

### Withdrawal score

Statistical analyses showed that the withdrawal score was different between groups (F_(4,45)_=14.8, p<0.001). Withdrawal score in morphine-dependent rats was more compared with healthy rats (p<0.0001). These results suggest that exposure to morphine was effective. On the other hand, there were no significant differences between dependent rats (p>0.05). Figure 3 shows withdrawal scores in the experimental groups.

**Figure 3.**
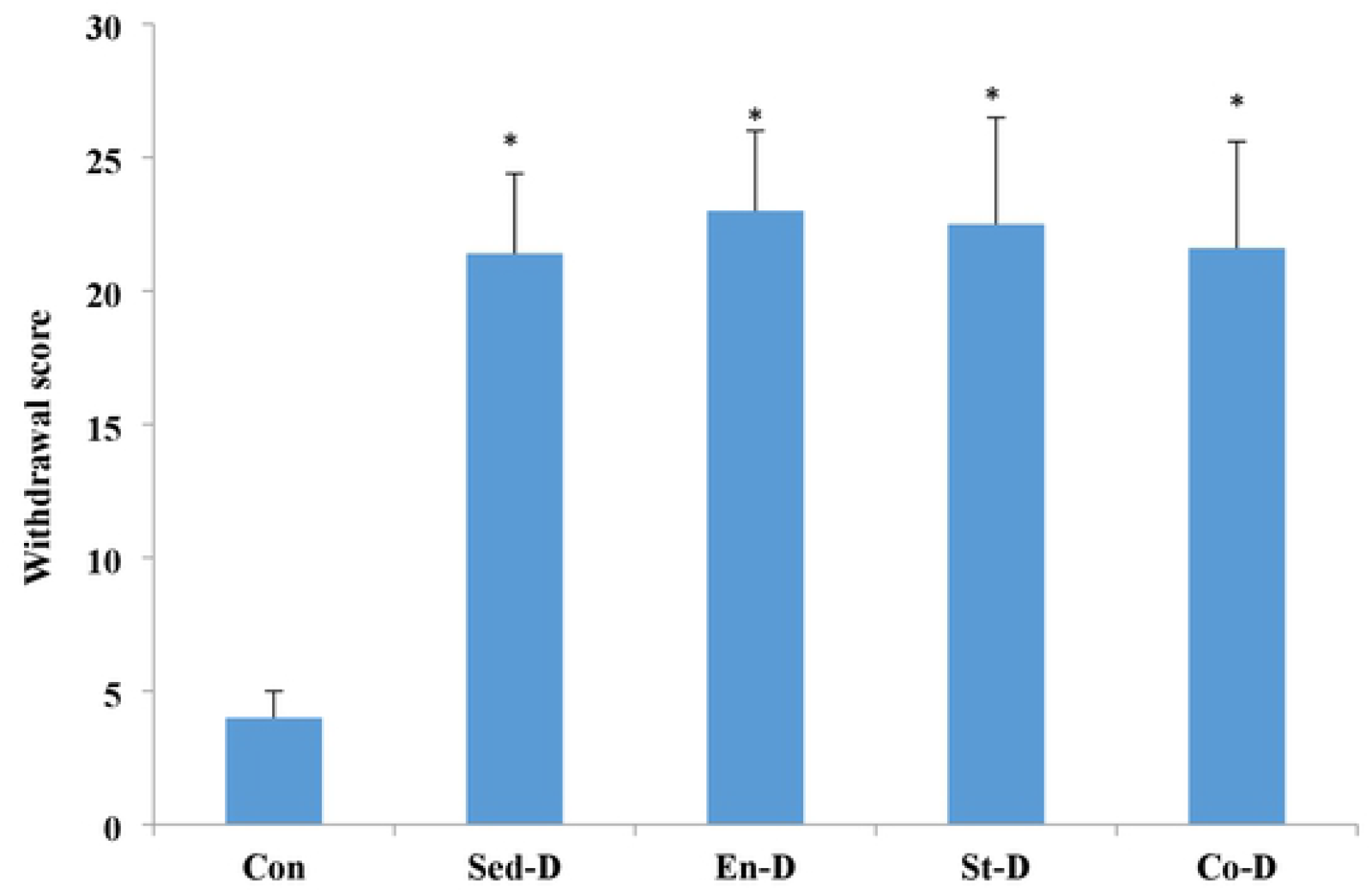
The scores of Gellert-Holztman in the experimental groups. Values are shown as mean ± SD. *P<0.05 vs.the control group.

### Measurement of evoked potentials

As shown in Figure 4, the changes in two components of each field potential, EPSP and PS, were recorded. The PS amplitude was evaluated from the peak of the initial positive deflection of the evoked potential to the peak of the next negative potential. Measurement of the fEPSP slope functional role was based on the slope of the line, by which the beginning of the first positive deflection of the evoked potential is connected to the peak of the second positive deflection, as we previously described [22, 32]. PP stimulation in the healthy and the morphine-dependent rats resulted in fEPSP slope as well as PS amplitude in the DG granular cells (Fig. 4).

**Figure 4.**
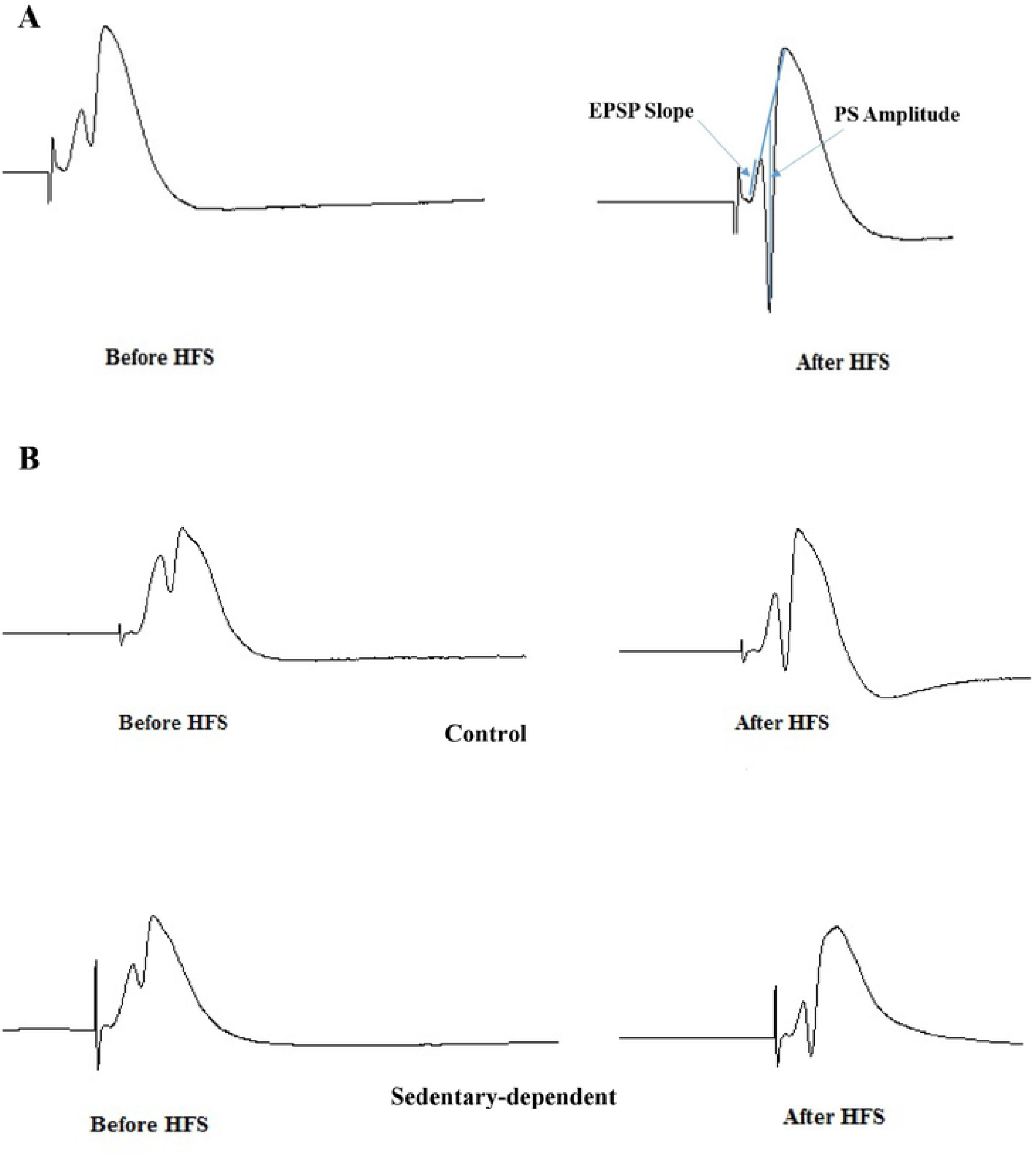
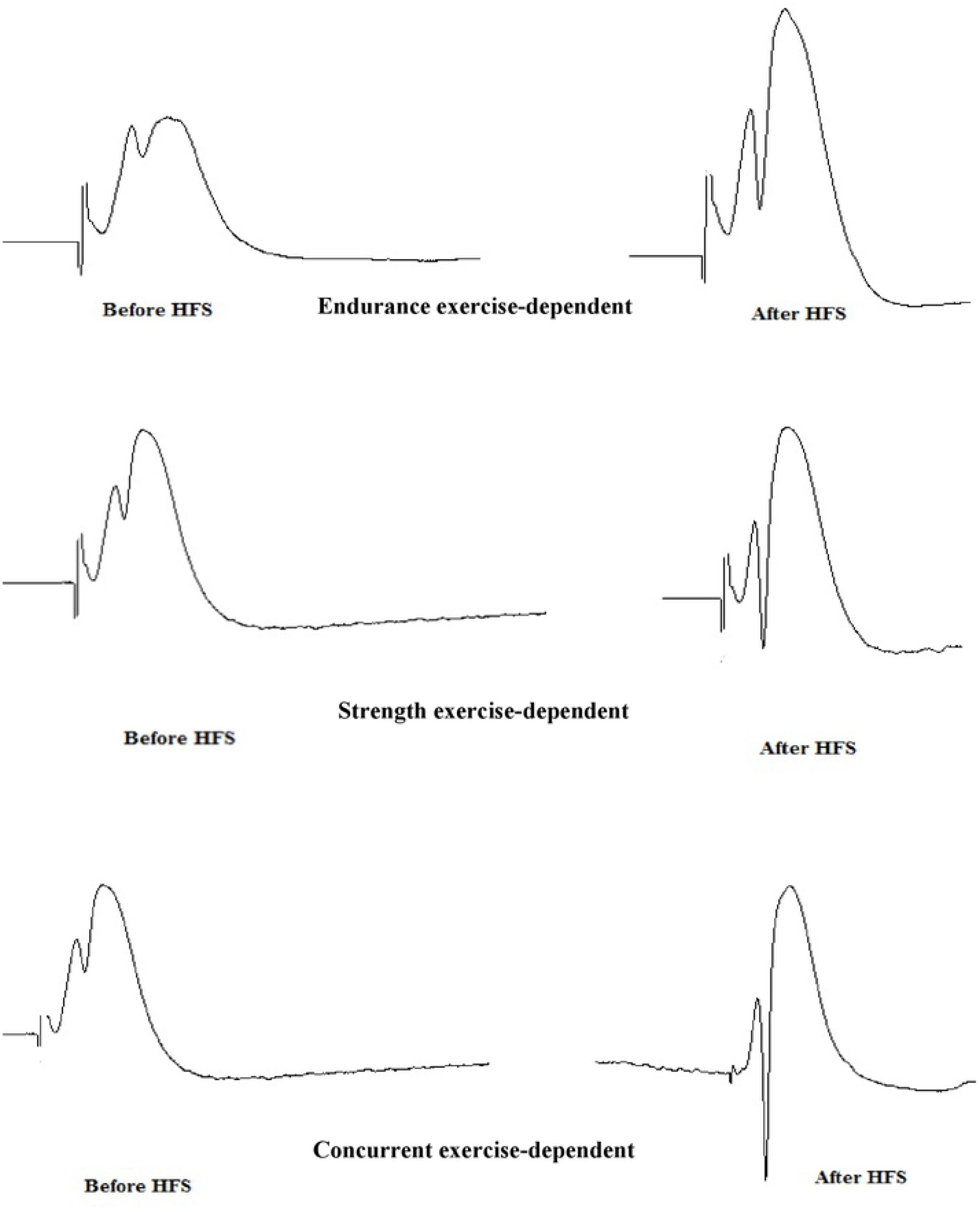
The excitatory postsynaptic potential (EPSP) slope and population spike (PS) amplitude of representative sample traces of field potential were obtained in the lateral perforant pathway-dentate gyrus (PP-DG) synapses of the control group. The arrows represent PSs and the slope of the EPSP (A). Representative sample traces of the evoked field potential in the DG, recorded before and following high-frequency stimulation (HFS) in all groups (B).

### HFS impact on the EPSP slope

Different exercise training effects on the HFS-induced EPSP slope and PS amplitude in the PP-DG region of the hippocampus in the healthy and the morphine-dependent rats are shown in Figures 5 and 6. The results of mixed-plot factorial repeated-measures ANOVA for EPSP slope indicated that there was a significant difference in terms of time (F _(3,135)_ =327.5, p<0.0001) and also a significant interaction was found between groups and time (F_(12,135)_=16.4, p<0.0001; Figure 5). Based on the findings, there was a significant difference among the experimental groups (F _(4,45)_=32.3, p<0.0001). In all groups, the highest increase in the HFS-induced EPSP slope was seen 5 min after HSF. The intragroup comparison demonstrated no significant difference in the EPSP in the Sed-D and control groups. In addition, no significant difference was observed in EPSP levels in the En-D group compared with the control and the Sed-D groups. Also, strength and concurrent training strongly increased HFS-induced EPSP in the PP-DG region of the hippocampus than the control, Sed-D, and En-D groups (p<0.0001). These results showed that concurrent and strength training, unlike aerobic training, can increase synaptic transmission and synaptic plasticity in the DG granular cells.

**Figure 5.**
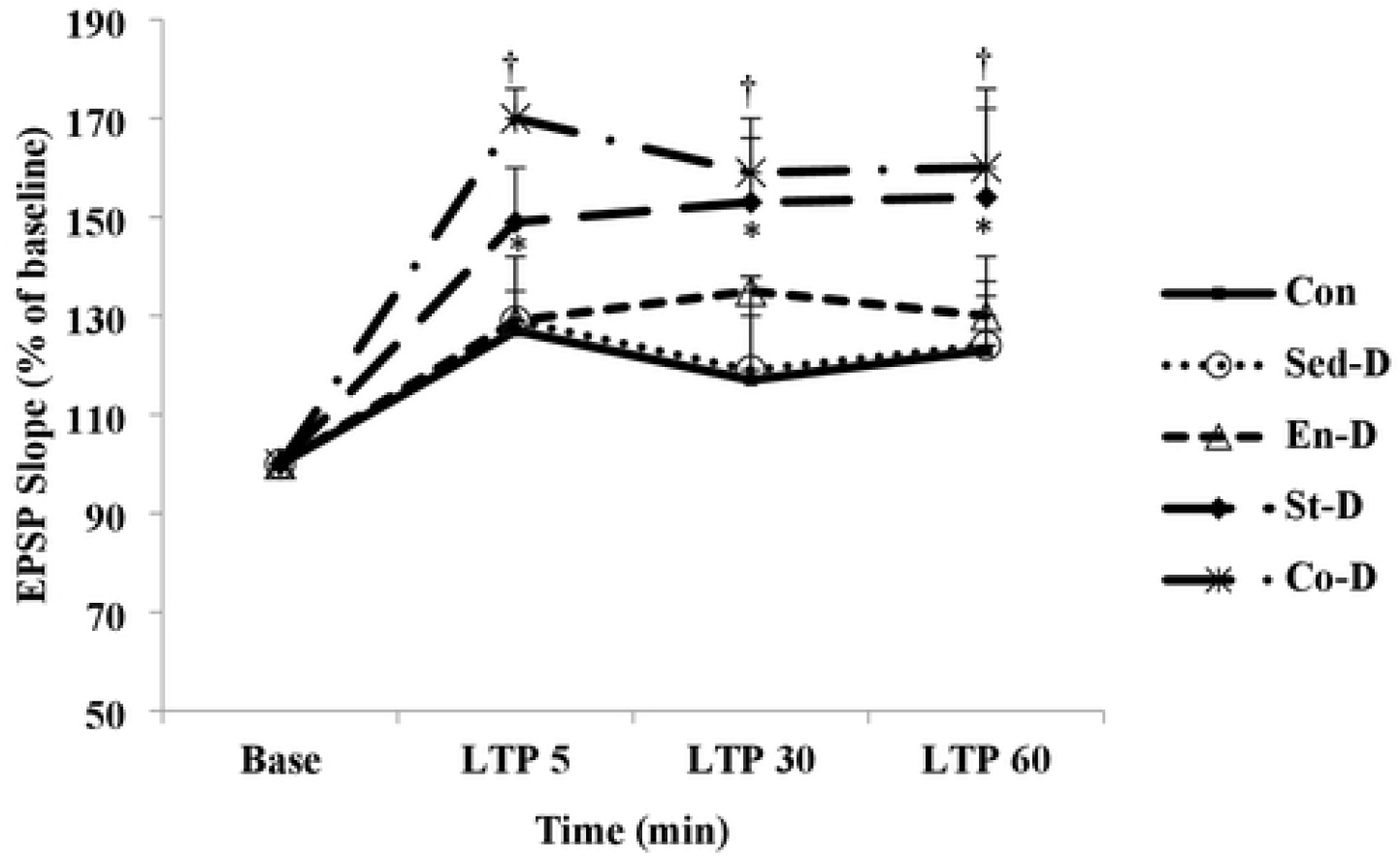
Time-dependent changes of the long-term potentiation (LTP) of the excitatory postsynaptic potential (EPSP) slope in granular cell synapses of the dentate gyrus (DG) after perforant path stimulation following a high-frequency stimulation (HFS). Values are shown as mean ± SEM % of the baseline. † P < 0.05 (Co-D group vs. Con, Sed-D, and En-D groups),* P < 0.05 (St-D group vs. Con, Sed-D, and En-D groups). Co-D: concurrent training-dependent group, St-D, strength training-dependent group, Sed-D: sedentary morphine-dependent group, Con: control group, En-D: endurance training-dependent group.

**Figure 6.**
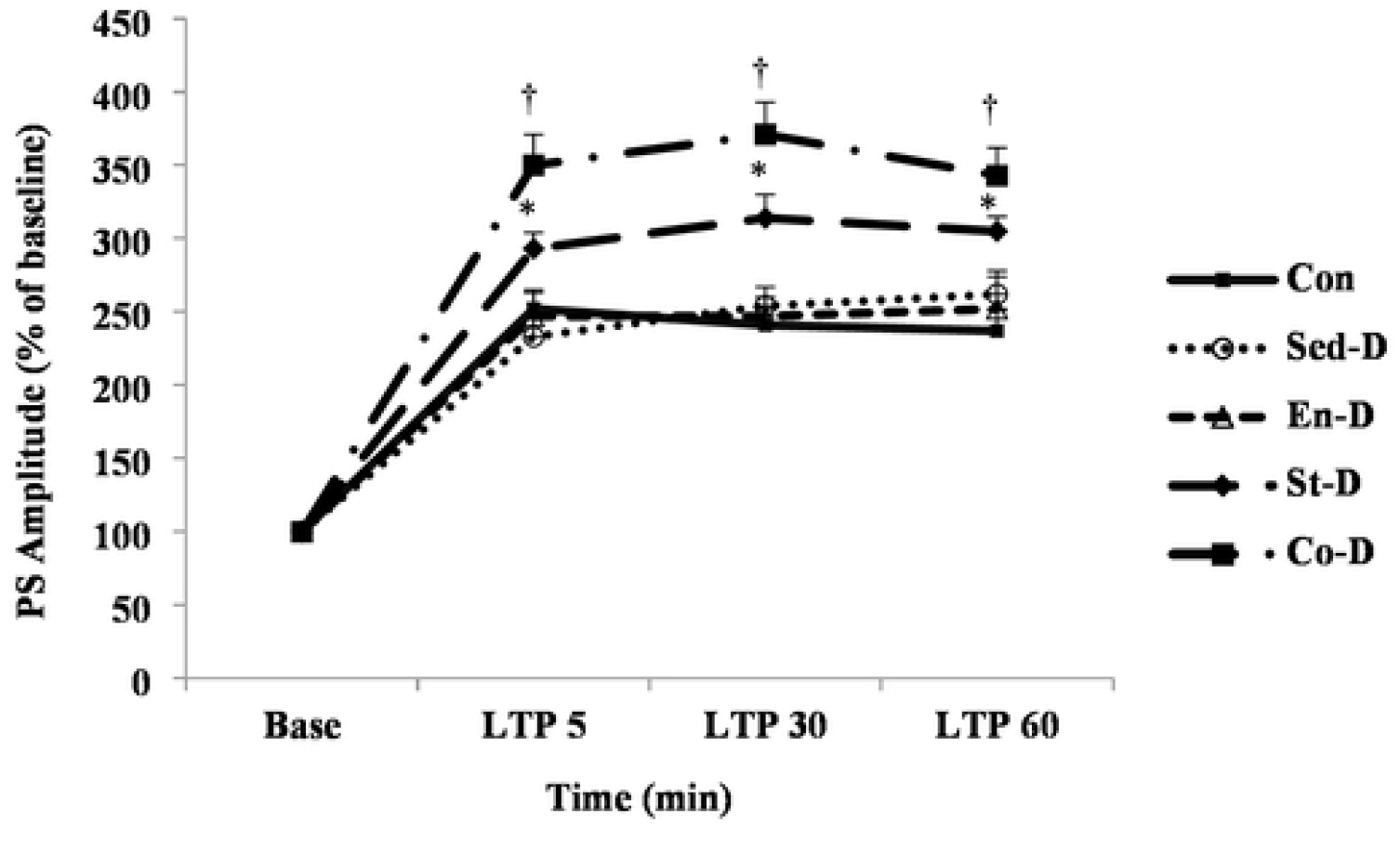
Time-dependent changes of the long-term potentiation (LTP) of the population spike (PS) amplitudes in dentate gyrus (DG) granular cell synapses after perforant path stimulation following a high-frequency stimulation (HFS). Values are shown as mean ± SEM % of the baseline. † P < 0.05 (Co-D vs. Con, Sed-D, En-D, and St-D groups),* P < 0.05 (St-D group vs. Con, Sed-D, En-D, and Co-D groups). Co-D: concurrent training-dependent group, St-D, strength training-dependent group, Sed-D: sedentary morphine-dependent group, Con: control group, En-D: endurance training-dependent group.

### HFS impact on the PS amplitude

As shown in Figure 6, statistical analysis on PS amplitudes showed significant effects of time (F_(3,135)_=627.5, p<0.0001) and also the group and time were found to be significantly correlated (F_(12,135)_=10.2, p<0.0001). The LTP of PS amplitudes in DG granular cell synapses in responses to PP stimulation following an HFS was different between the experimental groups (F _(4,45)_=17.7, p=0.0001). Also, 5 min after HFS, enhancement in the PS amplitude was greatest in all experimental groups. The changes in PS amplitudes were higher in the concurrent training group. Based on the results of post hoc analysis, chronic morphine addiction had no effects on PS amplitude in the hippocampus. These results showed that chronic morphine addiction had no effect on the magnitude of LTP in the CA1 region of hippocampal slices compared with healthy rats. On the other hand, no significant difference was found in the PS amplitude in the En-D group in comparison with the control and Sed-D groups. Accordingly, strength training significantly increased PS amplitude than that of the control, Sed-D, and En-D groups (p<0.01). However, PS amplitudes in the rats subjected to the concurrent training were higher compared with the other experimental groups (p<0.01). These results showed that concurrent training is more effective in the promotion of excitatory synapses efficiency.

### NeuroD-positive cell numbers

As shown in Figure 7-A-F, the number of NeuroD-positive cells, as an indicator for newborn immature neurons, was different among the experimental groups (F _(4, 20)_=14.8, P=0.0001). In this regard, a significant reduction was found in the Sed-D rats than the healthy group (p=0.0001). For each animal, the average number of dark brown cells representing the expression of NeuroD-was obtained by counting 5 serial sections. The number of NeuroD1-expressing cells per mm^2^ in healthy rats was 22±1 and in the Sed-D group was 13±1. This result showed that chronic opiate exposure could significantly decrease neurogenesis (p<0.0001). In this regard, a significant increase was observed in the number of NeuroD-positive cells after 10 weeks of concurrent training compared with the Sed-D group (p=0.002). Endurance and strength training increased NeuroD-positive cell numbers than the Sed-D group, but it was not significant (p=0.06 and p=0.29, respectively). These results indicated that concurrent training unlike aerobic and strength training promotes neurogenesis. The obtained results showed that 10 weeks of concurrent training attenuated deficit in neurogenesis due to morphine addiction by increasing the number of NeuroD-positive cells in the hippocampus.

**Figure 7.**
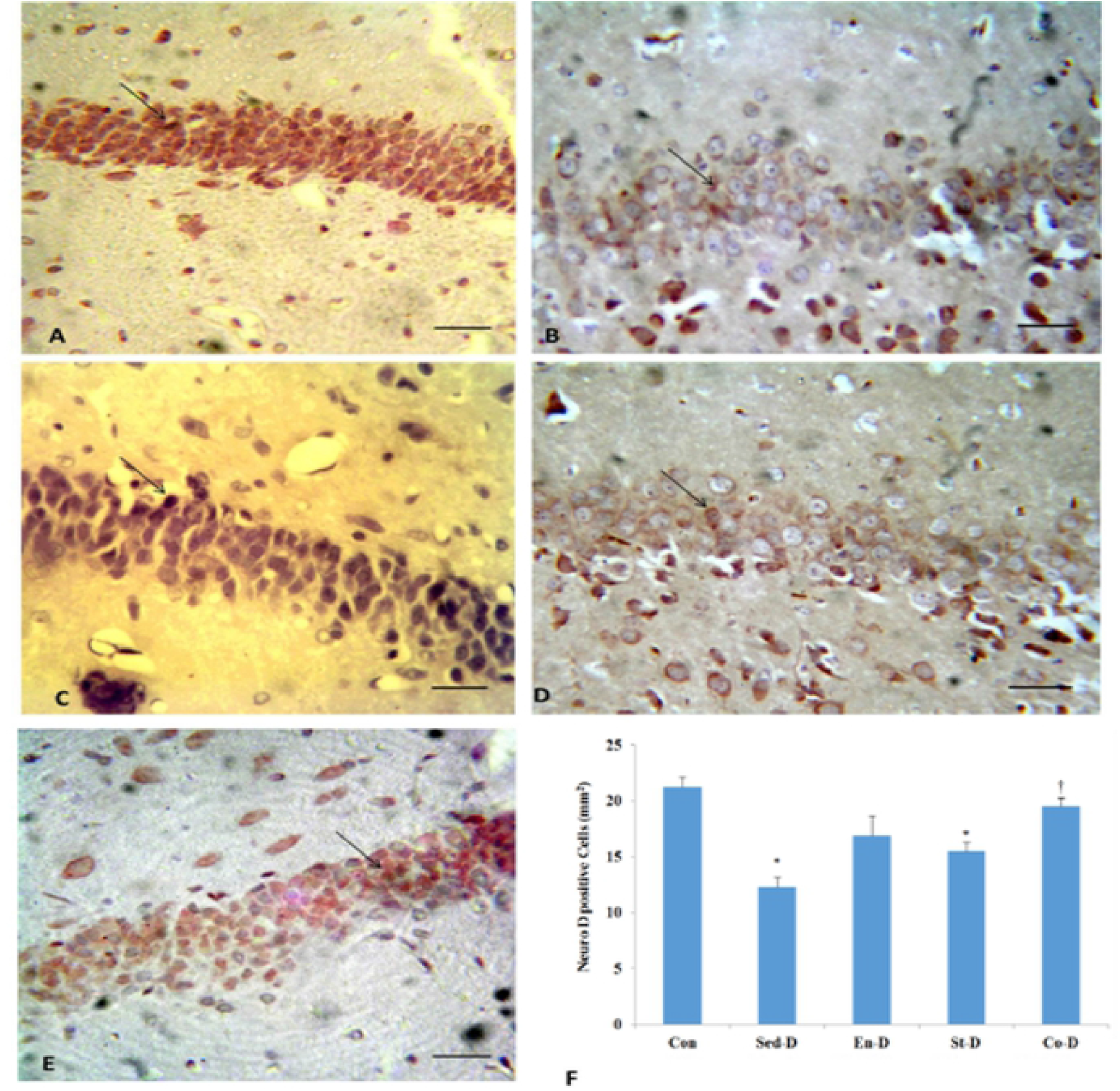
The NeuroD-positive cell numbers (mm2) in the rats’ dentate gyrus (DG) of the control (Con), sedentary morphine-dependent (Sed-D), endurance training-dependent (En-D), strength training-dependent (St-D), and concurrent training-dependent (Con-D) groups. The graph indicates the comparison between the NeuroD-positive cells in the experimental groups. The number of dark brown cells expressing NeuroD was counted. Arrows show NeuroD-positive cells (400×). The scale bar was 10 µm. Values are shown as mean ±SEM. *P < 0.05 vs. the Con group. † P < 0.05 vs. Sed-D group.

## Discussion

This research aimed at investigating the possible various effects of different exercise modes on LTP in the PP-DG granule cell synapse of the morphine-dependent rats in vivo. Briefly, according to the findings, although morphine addiction reduced NeuroD expression level, it had no effects on EPSP slope and SP amplitude, as indices of enhanced synaptic efficacy in DG granular cell synapses in responses to PP stimulation following an HFS. These results showed that morphine addiction did not lead to maladaptive changes in hippocampal plasticity.

The effect of morphine on LTP has been a controversial issue among researchers. Although it has been described with ameliorating effects [33], in other studies, it has been indicated with destructive effects [6, 10]. The chronic morphine administration in rats resulted in a significant decrease in the capacity of LTP in the CA1 hippocampal synapses through the drug withdrawal stage (from approximately 190% in the control to approximately 120%) [6]. It has been shown that the chronic administration of morphine (10 mg/kg at 12 h intervals for 10 days) caused an increase in the average basal EPSP and augmented LTP of the PS [33]. Salmanzadeh et al. indicated no changes in the fEPSP derived from dependent samples than the control group. This funding is in line with our results. Besides, the LTP of fEPSP in the CA1 area showed no difference after chronic morphine injection [34]. These contradictory results can be caused by the differences in the protocols of the studies. Moreover, previous studies have shown that the tetanus-induced synaptic plasticity may include various magnitudes or directions at any time point following withdrawal [10].

Exercise training promotes brain function, but its underlying neural mechanism has not yet been found [33]. We found that the EPSP slope and PS amplitude following HFS increased in the concurrent and strength training groups than the control group, which is in line with the findings of previous studies indicating that exercise training can revert the memory impairments induced by morphine [15, 17]. Therefore, the findings of our study suggested that exercise training can ameliorate cognitive function impairments caused by morphine dependence through LTP promotion.

In agreement with previous studies, our findings showed that concurrent training can promote synaptic plasticity in the hippocampal DG by increased NeuroD expression level. In this respect, it has also been indicated that exercise training can reduce apoptosis as well as stress oxidative and promote the expression of neurotrophic factors, cell proliferation, neurogenesis, and astrocytic plasticity [33, 35]. In addition, exercise training facilitates the induction of LTP by lowering the LTP induction threshold [33]. Regular exercise training prevents the decrement of the hippocampal DG neuron numbers induced by the injection of morphine through a brain-derived neurotrophic factor (BDNF)-mediated mechanism [36].

The most remarkable achievement of our study was that different exercise training programs have different effects on the hippocampus synaptic plasticity. Our results showed that unlike endurance training, strength, and concurrent training significantly increased LTP. It should be noted that different types of exercise can cause various impacts on brain function [37]. The effect of short-term aerobic exercise training (continuous and interval) on synaptic plasticity has been studied in a study at two different volumes. The results have demonstrated that there was an increase in the LTP calculated in the CA1 region only in animals trained with the interval 50% volume exercise compared with the sedentary group [38]. However, the exact causes of such changes are not clear. According to our findings, different impacts of various training modes on neuronal development can be considered as the main reason for such differences in LTP induction.

Our findings showed that the different effects of exercise modes on LTP were due to differences in the magnitude of the effect on neurogenesis. EPSP slope and PS amplitude 60 min after HFS in the Co-D group were 30% and 31% higher compared with the Sed-D group. These changes in percentage in response to concurrent training were higher compared with those observed in the strength (EPSP 23%, PS 16%) and endurance (EPSP 5%, PS -3%) training. However, the magnitude of neurogenesis promotion in the concurrent group was more compared with the endurance and strength groups (30%, 15%, and 92% promotion in the number of NeuroD-positive cells in response to endurance, strength, and concurrent training, respectively than the Sed-D group). Therefore, the effect of concurrent training on synaptic plasticity is dependent on neurogenesis amelioration.

Several complex mechanisms have been proposed for such differences, including differences in the density and/or distribution of NMDA receptors, mu-receptors, other kinds of glutamate receptors, intracellular Ca^2+^ concentration, and calmodulin-binding protein in the CA1 and DG areas. Furthermore, various exercise training types can protect the nervous system by the reduced oxidative stress as well as the improved antioxidant system and brain plasticity [39]. It has been reported that different training loads have remarkable influences on the cognition, emotion, and mental status of rats, and can affect the mRNA and protein expressions of NMDARs, PSD-95, and kinesin family member 17 (KIF-17) in rats [40]. In addition, cyclooxygenase (COX)-2 has a pivotal role in long-term synaptic plasticity modulation in the hippocampal PP-DG synapses [41]. Kruger et al. recently reported that the COX-2 expression induction is more obvious following strength training [42].

In summary, concurrent training prevents morphine-induced neurodegeneration via a rise in neurogenesis in the hippocampus DG of rats resulting in synaptic plasticity enhancement. Therefore, exercise training may be an appropriate novel candidate for treating opioid addiction.

## Acknowledgments

The staff of the Neurophysiology Research Center is appreciated for their cooperation.

## Author Contributions

Conceptualization: Iraj Salehi, Alireza Komaki.

Data curation: Kamal Ranjbar, Ebrahim Zarrinkalam, Masoumeh Taheri.

Formal analysis: Sara Soleimani Asl, Alireza Komaki.

Funding acquisition: Iraj Salehi, Alireza Komaki.

Investigation: Kamal Ranjbar.

Methodology: Sara Soleimani Asl, Alireza Komaki.

Project administration: Iraj Salehi, Alireza Komaki.

Software: Ebrahim Zarrinkalam, Masoumeh Taheri.

Supervision: Alireza Komaki, Iraj Salehi.

Writing – original draft: Kamal Ranjbar.

Writing – review & editing: Ebrahim Zarrinkalam, Iraj Salehi, Alireza Komaki.

